# Targeted Enrichment of Large Gene Families for Phylogenetic Inference: Phylogeny and Molecular Evolution of Photosynthesis Genes in the Portullugo (Caryophyllales)

**DOI:** 10.1101/145995

**Authors:** Abigail J. Moore, Jurriaan M. de Vos, Lillian P. Hancock, Eric Goolsby, Erika J. Edwards

## Abstract

Hybrid enrichment is an increasingly popular approach for obtaining hundreds of loci for phylogenetic analysis across many taxa quickly and cheaply. The genes targeted for sequencing are typically single-copy loci, which facilitate a more straightforward sequence assembly and homology assignment process. However, single copy loci are relatively uncommon elements of most genomes, and as such may provide a biased evolutionary history. Furthermore, this approach limits the inclusion of most genes of functional interest, which often belong to multi-gene families. Here we demonstrate the feasibility of including large gene families in hybrid enrichment protocols for phylogeny reconstruction and subsequent analyses of molecular evolution, using a new set of bait sequences designed for the “portullugo” (Caryophyllales), a moderately sized lineage of flowering plants (~2200 species) that includes the cacti and harbors many evolutionary transitions to C_4_ and CAM photosynthesis. Including multi-gene families allowed us to simultaneously infer a robust phylogeny and construct a dense sampling of sequences for a major enzyme of C_4_ and CAM photosynthesis, which revealed the accumulation of adaptive amino acid substitutions associated with C_4_ and CAM origins in particular paralogs. Our final set of matrices for phylogenetic analyses included 75–218 loci across 74 taxa, with ~50% matrix completeness across datasets. Phylogenetic resolution was greatly improved across the tree, at both shallow and deep levels. Concatenation and coalescent-based approaches both resolve with strong support the sister lineage of the cacti: Anacampserotaceae + Portulacaceae, two lineages of mostly diminutive succulent herbs of warm, arid regions. In spite of this congruence, BUCKy concordance analyses demonstrated strong and conflicting signals across gene trees for the resolution of the sister group of the cacti. Our results add to the growing number of examples illustrating the complexity of phylogenetic signals in genomic-scale data.

---

Next-generation sequencing has revolutionized the field of phylogenetics, and there are now many approaches available to efficiently collect genome-scale data for a large number of taxa. In one way or another, they all involve downsampling the genome as a means to simultaneously sequence homologous genomic regions across multiple species. Transcriptome analysis was among the first approaches (Dunn et al. 2008; Jiao et al. 2011; Wickett et al. 2014), and this remains an effective method, but typically fresh or flash-frozen tissues must be used for RNA extraction. Many researchers have large and invaluable collections of stored genomic DNA collected over years of fieldwork that must remain relevant. More recently, approaches such as RAD-seq, genome skimming, and hybrid enrichment have been adopted as effective means of sub-sampling the genome to enable development of very large datasets (1,000s of loci) across large numbers of individuals with multiplexed sequencing (McCormack et al. 2013). For deeper phylogenetic problems spanning larger clades, hybrid enrichment is emerging as the method of choice (Faircloth et al. 2012; Lemmon et al. 2012; de Sousa et al. 2014; Mandel et al. 2015; Schmickl et al. 2015; 2016).

Hybrid enrichment studies tend to limit their scope to ‘single-copy loci’ (SCL), that is, genes that do not appear to have maintained multiple paralogs within a genome after a gene duplication. Targeting SCL has obvious appeal, as it facilitates straightforward contig assembly and reduces the risk of constructing erroneous gene trees due to incorrect orthology assignment. However, the number of SCL in a genome is relatively small, and especially in plants, they appear to be somewhat unusual. As all extant flowering plants have undergone multiple rounds of whole genome duplication (De Bodt et al. 2005; Jiao et al. 2011; Renny-Byfield and Wendel 2014; Soltis et al. 2015), SCL are likely under strong selection to lose additional gene copies after undergoing duplication (Freeling 2009; de Smet et al. 2013). If gene loss happens very quickly post duplication (i.e., prior to subsequent speciation events), these loci would be especially useful for phylogenetics; if, on the other hand, gene loss is more protracted, these loci could instead be especially problematic. Genome-wide estimates suggest that the rate of duplication is quite high (0.01/gene/Ma) and subsequent loss is relatively slow, with the average half-life of a duplicate gene estimated at ~4 Myr (Lynch and Conery 2000). It is at least worth considering that purported SCL may be susceptible to “hidden” paralogy issues, due to differential loss of duplicates over longer periods of time (Martin and Burg 2002; Álvarez and Wendel 2003).

An additional limitation of constraining analyses to SCL is the necessary omission of genes of potential interest for other sorts of evolutionary or functional studies, independent of their utility in phylogenetic inference. Beyond the primary goal of generating data for phylogenetic inference, hybrid enrichment offers an unparalleled potential to affordably and efficiently build large comparative datasets of important functional genes, enabling molecular evolution analyses of a scope not seen before. Because the substrate of hybrid enrichment is whole genomic DNA, rather than transcriptomic data of expressed genes, there is also the potential to isolate additional copies of genes that were not expressed at the time of tissue collection, providing a more complete picture on the evolutionary dynamics of gene duplication and function/loss of function. There are several methodological challenges to unlocking this potential, including: 1) designing baits that can target multiple members of large gene families across disparate groups of taxa, 2) accurately joining fragmented contigs that belong to the same paralog within individuals together into a single non-chimeric locus, and 3) confident assignment of loci to their correct orthologs across species. Each of these tasks is difficult but, we demonstrate, not insurmountable.

We present a first attempt to include multi-gene families in a hybrid enrichment study of the “Portullugo” (Caryophyllales) (sensu Edwards and Ogburn 2012), a diverse clade of ~2200 species of flowering plants with a worldwide distribution (Fig. 1). The clade includes nine major lineages, is most commonly found in warm and arid or semi-arid environments, and includes such charismatic succulents as the cacti of the New World and the Didiereaceae of Madagascar. The portullugo has received a fair amount of phylogenetic attention over the decades (e.g., Hershkovitz and Zimmer 2000; Applequist and Wallace 2001; Nyffeler and Eggli 2010; Ocampo and Columbus 2010; Ogburn and Edwards 2015), yet relationships among many of its major lineages remain stubbornly unresolved; one particularly recalcitrant region is the relationship between the cacti, *Portulaca*, and Anacampserotaceae.

**Figure 1.**
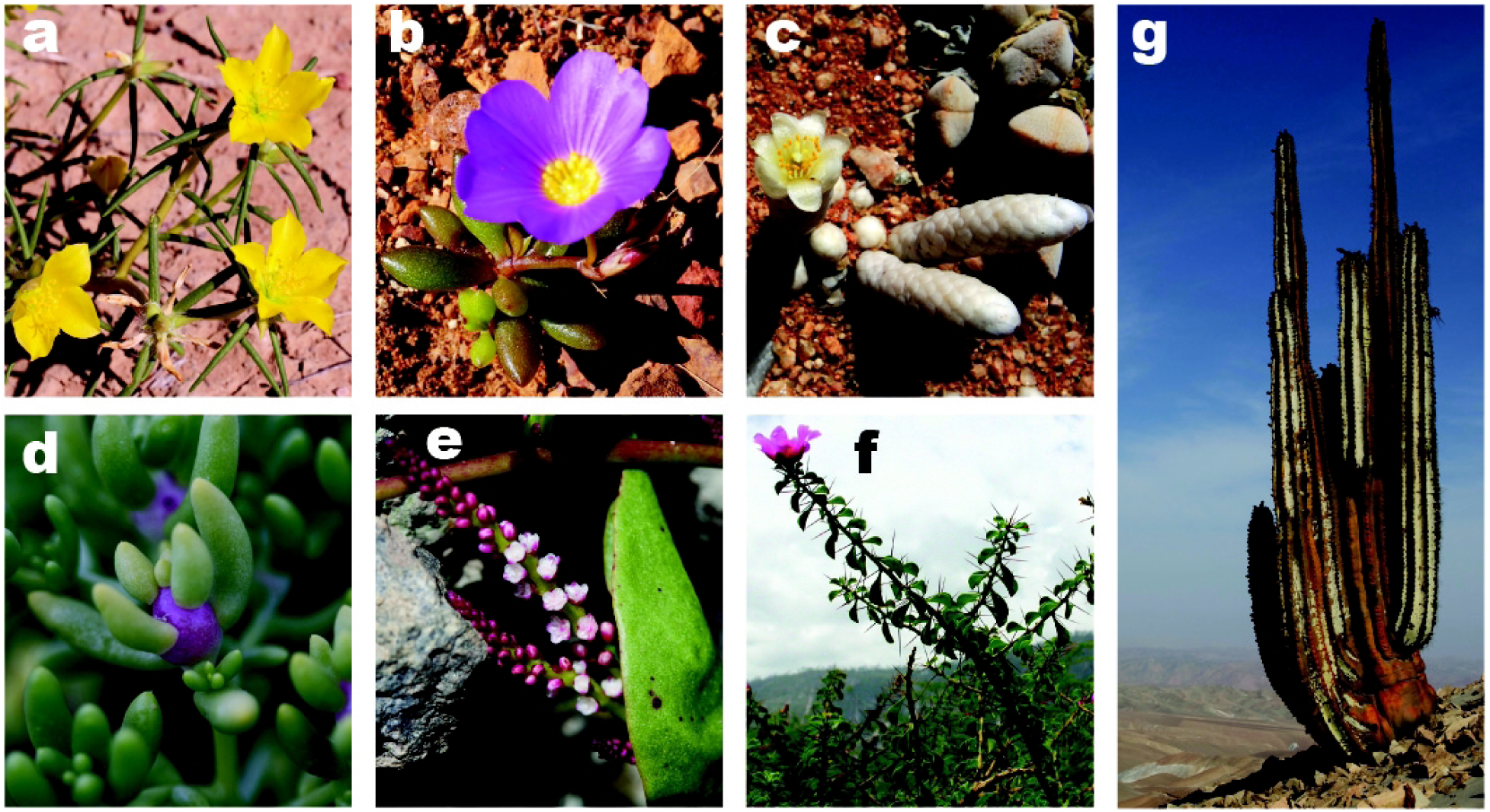
Species representatives across Portulacineae families. A) *Portulaca aff. filsonii* B) *Calandrinia hortiorum* C) *Anacampseros papyracea* D) *Halophytum ameghinoi* E) *Anredera diffusa*, F) *Pereskia portulacifolia*, G) *Neoraimondia arequipensis*.

Portullugo also harbors multiple origins of two plant metabolic pathways: C_4_ and CAM photosynthesis, both complex syndromes that employ a shared set of enzymes to increase internal plant CO_2_ concentrations and improve photosynthetic efficiency (Edwards and Ogburn 2012). We are especially interested in the molecular evolution of genes coding for the major C_3_, C_4_, and CAM photosynthesis enzymes during evolutionary transitions between these metabolic pathways, and included 19 major photosynthesis gene families in our hybrid enrichment design. Phylogenetic analyses of our data resolve many outstanding issues in portullugo phylogeny, and we also present the utility of our dataset for analyzing adaptive protein sequence evolution, with a preliminary analysis of the PEP Carboxylase gene family. In both C_4_ and CAM photosynthesis, PEP Carboxylase is the enzyme recruited to first fix atmospheric CO_2_ in leaves, where it is temporarily stored as a 4-carbon acid and later decarboxylated in the presence of the Calvin cycle. The enzyme is a critical component of both pathways, and previous work has demonstrated convergent evolution of multiple amino acid residues associated with both C_4_ and CAM origins (e.g., Christin et al. 2007; 2014).

## Materials and Methods

### Terminology

We use the term paralog to describe gene copies that diverged from one another in a duplication event, hence multiple paralogs can be present in a single individual. In contrast, ortholog is used when referring to a set of homologous genes that originated via speciation events. Depending on the context, a single gene can therefore be included and discussed in the context of a paralog group or an ortholog group. In the context of phylogenetic inference involving all sequenced genes, we refer to all orthologs from all of the various gene families as loci.

### Data Availability

All scripts are available in a public repository. One folder contains the analysis pipeline (https://github.com/abigail-Moore/baits-analysis) and a second folder contains scripts for bait design and gene tree/species tree analysis (https://github.com/abigail-Moore/baits-suppl_scripts).

### Probe Design

Probes for targeted enrichment were designed for use across the portullugo based on analyses of eight previously sequenced transcriptomes from the Portulacineae (Christin et al. 2014, 2015; Anacampserotaceae: *Anacampseros filamentosa*; Cactaceae: *Echinocereus pectinatus, Nopalea cochenillifera, Pereskia bleo, Pereskia grandifolia, Pereskia lychnidiflora*; Portulacaceae: *Portulaca oleracea*; and Talinaceae: *Talinum portulacifolium*) and four from its sister group Molluginaceae (Matasci et al. 2014; *Hypertelis cerviana* (called *M. cerviana* in 1KP), *Mollugo verticillata, Paramollugo nudicaulis* (called *M. nudicaulis* in 1KP), and *Trigastrotheca pentaphylla* (called *M. pentaphylla* in 1KP)). MyBaits probes were designed from two sets of genes: 19 gene families that were known to be important in CAM and C_4_ photosynthesis, and 52 other low- or single-copy nuclear genes (Table S1; MYcroarray, Ann Arbor, MI, USA).

Sequences for photosynthesis-related genes were taken from the alignments from Christin et al. (2014, 2015), which included the transcriptomic data, sequences from GenBank, and individual loci from other members of the portullugo clade. Gene family identities for the remaining genes in the portullugo transcriptomes were assigned by blasting (BLASTN 2.2.25, default settings; Altschul et al. 1990) them against sets of orthologous sequences of known identity from six model plants (Ensembl database; Kersey et al. 2016; http://plants.ensembl.org/, accessed 4 Dec. 2013). Similarly, we also assigned gene family identities to genes from five additional Caryophyllales transcriptomes, which had previously been sequenced (*Amaranthus hypochondriacus*, Amaranthaceae; *Boerhavia coccinea*, Nyctaginaceae; *Mesembryanthemum crystallinum*, Aizoaceae; *Trianthema portulacastrum*, Aizoaceae; Christin et al. 2015; *Beta vulgaris*, Amaranthaceae, Dohm et al. 2014) to be able to include them in subsequent analyses. Further details of probe design are provided in the Supplementary Methods.

### Taxon Sampling

Sixty portullugo individuals were sequenced (Supplemental Table 2), including multiple representatives of all major lineages (with the exception of the monotypic Halophytaceae, which was represented by *Halophytum ameghinoi*), and relevant sequences from transcriptomes of two further species were added (*Pereskia bleo*, Cactaceae; *Portulaca oleracea*, Portulacaceae). Eleven outgroups were added by extracting the relevant sequences from the five non-portullugo, Caryophyllales transcriptomes and the six model plant genomes, for a total of 73 taxa.

### Molecular Sequencing

Leaf material was first extracted using the FastDNA Spin Kit (MP Biomedicals, Santa Ana, CA). After DNA extraction, samples were cleaned using a QIAquick PCR Cleanup Kit (Qiagen Inc., Valencia, CA), following the manufacturer’s protocol. DNA was fragmented using sonication and libraries were prepared using the NEBNext Ultra or NEBNext Ultra II DNA Library Prep Kits for Illumina (New England Biolabs, Ipswich, MA), including addition of inline barcodes (see supplementary methods for details). For hybridization with MyBaits probes, samples were combined into groups of 8 or 9 with approximately equal amounts of DNA for each sample, resulting in a total of 100–500 ng of DNA in 5.9 μl of buffer. A low stringency hybridization protocol was followed, because species used for bait design were sometimes distantly related to the species sequenced (Li et al. 2013). The remainder of the hybridization and cleanup protocol followed version 2 of the manufacturer’s protocol, except that the cleanup steps took place at 50°C instead of 65°C. Final quantification, combination, and sequencing of most samples were performed at the Brown University Genomics Core Facility on an Illumina HiSeq 2000 or 2500, to obtain 100-bp, paired end reads. Further details are given in the supplementary methods. Reads for each individual were submitted to the NCBI SRA (BioProject PRJNA387599, accession numbers in Supplemental Table 2).

### Methods Summary for Data Processing and Orthology Assignment

We designed a three-part bioinformatics pipeline to reconstruct gene sequences (Fig. 2). Part I aimed to extract all relevant reads for each gene family and then assemble them into contigs. Part II then constructed longer sequences from contigs and assigned them to particular paralogs within a gene family. Part III identified gene duplications within gene families, extracted phylogenetically useful sets of orthologs, and used them for phylogenetic analysis. The major steps in the pipeline are summarized below; details are given in the supplementary methods.

**Figure 2.**
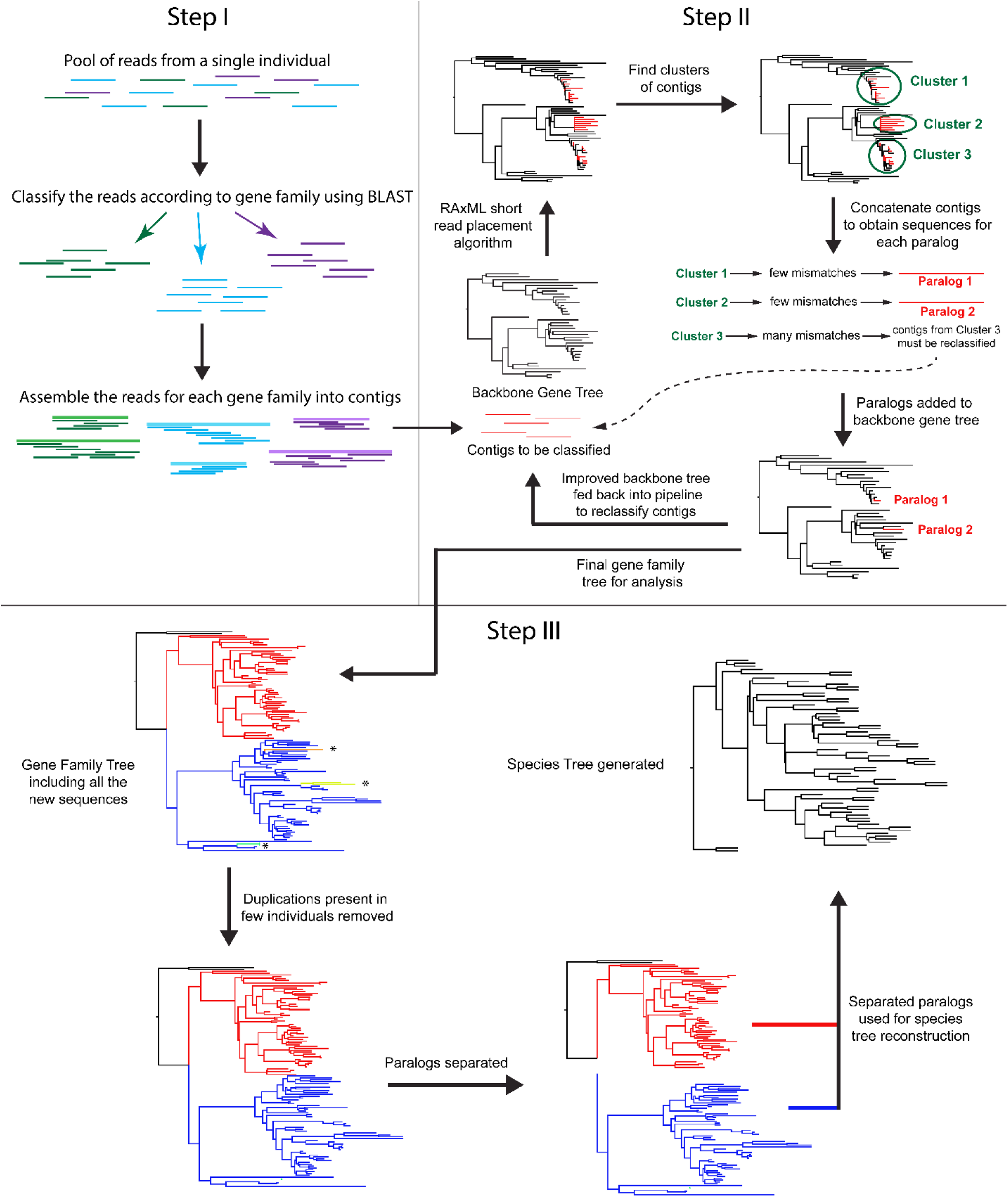
Assembly pipeline schematic. Part I extracts all relevant reads for each gene family and then assemble them into contigs. Part II constructs longer sequences from contigs and assigns them to particular paralogs within a gene family. Part III identifies gene duplications within gene families, isolates phylogenetically useful sets of orthologs, and uses them for phylogenetic analysis.

In part I, paired reads were classified into gene families using BLASTN version 2.2.29 (Altschul et al. 1990) and assembled into contigs. A read pair was assigned a gene family if either read matched (e-value < 10–^16^) the sequences used to design its baits. For each gene family, reads were then pooled among the individuals that belonged to each of the 9 major lineages, and SPAdes version 3.1.0 (Bankevich et al. 2012) was used to assemble them into 9 preliminary contigs. By using reads from different individuals and different species in the same assembly, we maximized contig number and lengths by also assembling chimeric contigs containing reads from multiple individuals; this step allowed us to pull significantly more reads into the pool for analysis. In the next step, a new BLAST database was created from the chimeric contigs and the sequences from which the baits were designed. The reads were then blasted to this larger database, again extracting both reads of a pair if either matched. For each individual and gene family, reads were assembled using SPAdes. Finally, the resulting contigs were blasted against the bait sequences to identify exons, and only exons were used for all subsequent analyses.

Part II of the pipeline identified the paralog that each contig from Part I belonged to, in order to combine contigs and maximize the sequence length for each paralog. This iterative process began with initial backbone alignments and trees for each gene family; these consisted of the sequences used to design the baits as well as the model plant and non-portullugo Caryophyllales sequences. In each iteration, all contigs for a gene family were first added to the backbone alignment using MAFFT version 7.017 (Katoh and Standley 2013) and then placed in the backbone gene family tree using the short-read classification algorithm in RAxML version 8.0.22 (option “-f v”; Berger 2011, Stamatakis 2014). These two steps yielded gene-family trees that contained one or several clusters of contigs. Each cluster was treated as a putative paralog and extracted for further testing. For each cluster and individual, contigs were combined into a consensus sequence (based on their positions in the backbone alignment) if the number of conflicting bases in overlapping contigs (e.g., due to presence of multiple alleles) was acceptably low. If a consensus sequence was successfully produced from a cluster of contigs, it was added to the backbone tree. If not, those contigs were analyzed again in the next iteration. After six iterations, some contigs could still not be combined into acceptable consensus sequences (e.g. due to recent gene duplications that were absent from the backbone tree). Here, a single contig per individual and paralog was selected.

Part III of the pipeline extracted paralogs as separate phylogenetic loci from the gene-family trees, by identifying the positions of gene duplications in comparison with a preliminary species tree, and used these loci to reconstruct gene trees. Part III was performed twice, first with a preliminary species tree constructed from three chloroplast loci (*matK, ndhF*, and *rbcL*) and the nuclear internal transcribed spacer (ITS) region, all recovered as off-target reads, and then with an updated species tree, reconstructed from the loci recovered from the pipeline. For each round of analysis, NOTUNG version 2.8.1.6 (Chen et al. 2000, Stolzer et al. 2012) was used to find gene duplications in the gene family trees, based on the given species tree. While the topology of the species tree was taken as given, poorly supported nodes (< 90% bootstrap) on the gene family trees were rearranged to correspond to the species tree, to minimize the impact of lack of support on paralog classification. Besides accounting for poorly-supported incongruences between the gene-family tree and species tree, we also employed a conservative strategy involving a variety of criteria to accept duplications. Most importantly, a duplication was accepted if the two sister groups subtended by a putative duplication contained at least one shared individual or two shared taxonomic families represented by different individuals. This strategy prevented us from accepting putative duplications that in fact represent incongruence between the topologies of the gene family tree and species tree. After inspection, at each node that subtended an accepted duplication, the smaller sister group was pruned off as a distinct locus, while the larger group was retained on the gene tree. (Note that after pruning, the larger group represents more than a single paralog, as it also contains the unduplicated sequences from the tree partition not affected by the focal duplication.) This strategy maximized the number of loci that contained all or most of the individuals, facilitating phylogenetic inference. We then calculated the number of individuals and number of major lineages present in each locus, and removed all sites with >90% missing data prior to analysis.

### Reconstruction of Species Trees and Estimating Gene Tree Congruence

To evaluate phylogenetic relations and branch support, we used concatenation and coalescent-based approaches on each of five data sets. We selected the following data sets, after dividing species into eleven taxonomic groups (the nine major clades recognized within the portullugo, and two additional paraphyletic groups for the outgroups, namely non-portullugo Caryophyllales and non-Caryophyllales; Table S3, Fig. S1): all loci that were present in two or more groups (g2 matrix, 218 loci, 42.7% missing loci, where “missing loci” are loci that were completely absent for certain individuals), all loci present in at least 5 taxonomic groups (g5 matrix, 163 loci, 27.7% missing loci), all loci present in at least 50% of individuals (i36 matrix, 136 loci, 20.6% missing loci), all loci present in at least 9 taxonomic groups (g9 matrix, 115 loci, 18.1% missing loci), and all loci present in at least 80% of individuals (i57 matrix, 75 loci, 10.1% missing loci). Concatenation analyses were performed in RAxML with 100 bootstrap replicates. Coalescent-based species trees were reconstructed using ASTRAL II version 4.10.2 (Mirarab and Warnow 2015, Erfan and Mirarab 2016) using gene trees from RaxML as input.

In addition, to evaluate genomic support for relations among major clades of portullugo, Bayesian concordance analysis was performed using BUCKy version 1.4.4 (Larget et al. 2010) based on the posterior distribution of gene trees from analyses in MrBayes 3.2 (Ronquist et al. 2012). BUCKy estimates the genomic support as a concordance factor for each relationship found across analyses of all individual loci (Ane et al. 2006; Baum 2007)̤ This way, groups of genes supporting the same topology are detected, while accounting for uncertainty in gene tree estimates. BUCKy thus alleviates the concern that methods to reconciliate gene trees, such as ASTRAL, may underestimate uncertainty of species tree (Leache & Rannala 2011), by highlighting genomic conflict. The implementation of BUCKy analyses requires identically named tips to be present in trees for all loci. In order to maximize the number of loci that could be simultaneously analyzed, taxa were renamed to their major lineage (outlined above), and all but one random exemplar per family was pruned from each sample of the posterior distribution of MrBayes trees. Although there is strong support for the monophyly of the major portullugo lineages (Nyffeler and Eggli 2010), our renaming and pruning approach does not require it to be so for each individual gene tree. Rather, the phylogenetic position of a family is averaged out over all probable positions, because a large number of renamed, pruned trees from each posterior distribution are input.

We conducted two sets of BUCKy analyses: one focusing on the position of Cactaceae within theACPT clade (Anacampserotaceae + Cactaceae + Portulacaceae + Talinaceae), and a broader Portulacineae-wide analysis, focusing on the remaining relationships after collapsing the ACPT clade to a single taxon. MrBayes analyses generated a posterior distribution of gene trees for each locus and consisted of two runs of 4,000,000 generations with default MCMCMC settings, sampling every 4,000 generations, employing a GTR substitution model with gamma-distributed rate variation across sites. After confirming the adequacy of these settings and excluding 25% of samples as burnin, runs were combined to a full posterior distribution of 1500 samples and subsequently thinned to 200 samples. The full posterior distribution was subjected to the renaming and thinning approach described above. For each locus, posterior probabilities of the monophyly of each lineage and their relationships to one another were scored from the thinned posterior distribution as the fraction of sampled trees that contained the node of interest. Analyses were conducted on all loci in which all focal lineages were present (ACPT: n=143; Portulacineae: n=132). For all analyses, we ran BUCKy using four runs of 100,000 generations each and computed genome-wide concordance factors (in which loci are interpreted as a random sample from the genome) of all possible relationships, as well as the posterior probability of each locus pair to support the same tree. All processing of MrBayes and BUCKy files was performed using custom R scripts.

### Molecular Evolution of PEP Carboxylase

Phylogenetic trees of the three major lineages of PEP Carboxylase in eudicots (*ppc-1E1, ppc-1E2*, and *ppc2*) were inferred using RAxML. Coding sequences were translated into amino acid sequences and numbered according to *Zea mays* sequence CAA33317 (Hudspeth and Grula 1989). Fourteen amino acid residues (466, 517, 531,572, 577, 579, 625, 637, 665, 733, 761, 780, 794, and 807) that were previously determined to be under positive selection in C_4_ grasses (Christin et al. 2007; Besnard et al. 2009), as well as position 890, which is associated with malate sensitivity (Paulus et al. 2013), were examined across the three major paralogs separately. Some residues could not be identified due to missing data or ambiguity. For these residues, marginal ancestral state reconstruction was performed using the *rerootingMethod* function in the R package phytools (Revell 2012) to determine the amino acid with the highest marginal probability.

## Results

### Sequence Coverage and Dataset Structure

We obtained between 682,702 and 13,008,046 reads per individual (mean 3,385,697 ± 2,953,383; Table S2). Percent enrichment, expressed as the percentage of read pairs yielding a blast hit to the bait design alignments, ranged from 0.26% to 12.32% across individuals, with a mean of 2.66 ± 1.81% after two rounds of blasting (0.17% to 6.20%, with a mean of 1.72 ± 0.98% after one round of blasting; Table S2) and did not differ between species closely related to individuals with transcriptomes (Cactaceae, *Portulaca, Mollugo*) and more distantly related species (2.73 ± 1.01% vs 2.63 ± 2.56%, N = 59, p = 0.40 for 1-tailed t-test, samples having unequal variance).

Part I of the analysis pipeline yielded a widely varying number of contigs per individual and gene family (mean of 15.6 ± 35.5, range of 0 (in numerous cases) to 1816 (PEPC genes in *Ullucus tuberosus*)). The total number of contigs per individual ranged from 500 (*Calandrinia lehmannii*) to 2576 (*Ullucus tuberosus*), with a mean of 1123 ± 451 (Table S2). Part II of the pipeline consolidated these contigs into longer sequences, and the total number of sequences per individual ranged from 62 (*Calandrinia lehmannii*) to 221 (*Alluaudia procera*), with a mean of 149 ± 28 (Table S2). The number of loci per individual per gene family was also variable (mean of 1.85 ± 1.61, range of 0 (in numerous cases) to 13 (*nadmdh* in *Alluaudia procera*)).

The pipeline yielded a total of 665 phylogenetic loci, with the number of individuals per locus ranging from 1 to 72 and the number of loci per gene family varying between 1 (i.e., putatively single-copy; 5 loci) and 34 (*nadmdh*). Taxon sampling across these loci was quite variable, with some loci being present in all major lineages, and others only being present in a single group (because they were paralogs due to a gene duplication near the tips). The mean sequence length per locus varied between 152 and 4075 bp (812 ± 616 bp). There was considerable variation in the number of loci per gene family, both between gene families and between individuals. Photosynthesis genes generally had many more paralogs than the non-photosynthesis genes, although there was variation in both groups of gene families (Fig. S2 for heatmaps showing the number of recovered per gene family for each individual; Fig. S3 for duplication numbers across all branches of the species tree).

### Phylogenetic Analyses

Most nodes were congruent and well supported across all analyses of all matrices (Fig. 3; Fig. S4, all remaining trees). The major differences are summarized in Table S5. Most of the conflict between analyses concerned relationships within the cacti, particularly the various species of *Pereskia*, and some of the relationships among closely related species of the Montiaceae. All nine of the major clades within the portullugo were well supported (>95% bootstrap) in both coalescent (ASTRAL) and concatenated (RAxML) species trees. The following larger clades were also always well supported: Anacampserotaceae and Portulacaceae as sister lineages; the clade comprised of Anacampserotaceae, Cactaceae, and Portulacaceae (ACP); ACP plus Talinaceae (ACPT); ACPT plus Didiereaceae; and the Portulacineae (portullugo without Molluginaceae). The analyses consistently recovered Montiaceae alone as sister to the seven remaining clades in the Portulacineae, and Basellaceae as sister to Halophytaceae, although with lower support in both cases.

**Figure 3.**
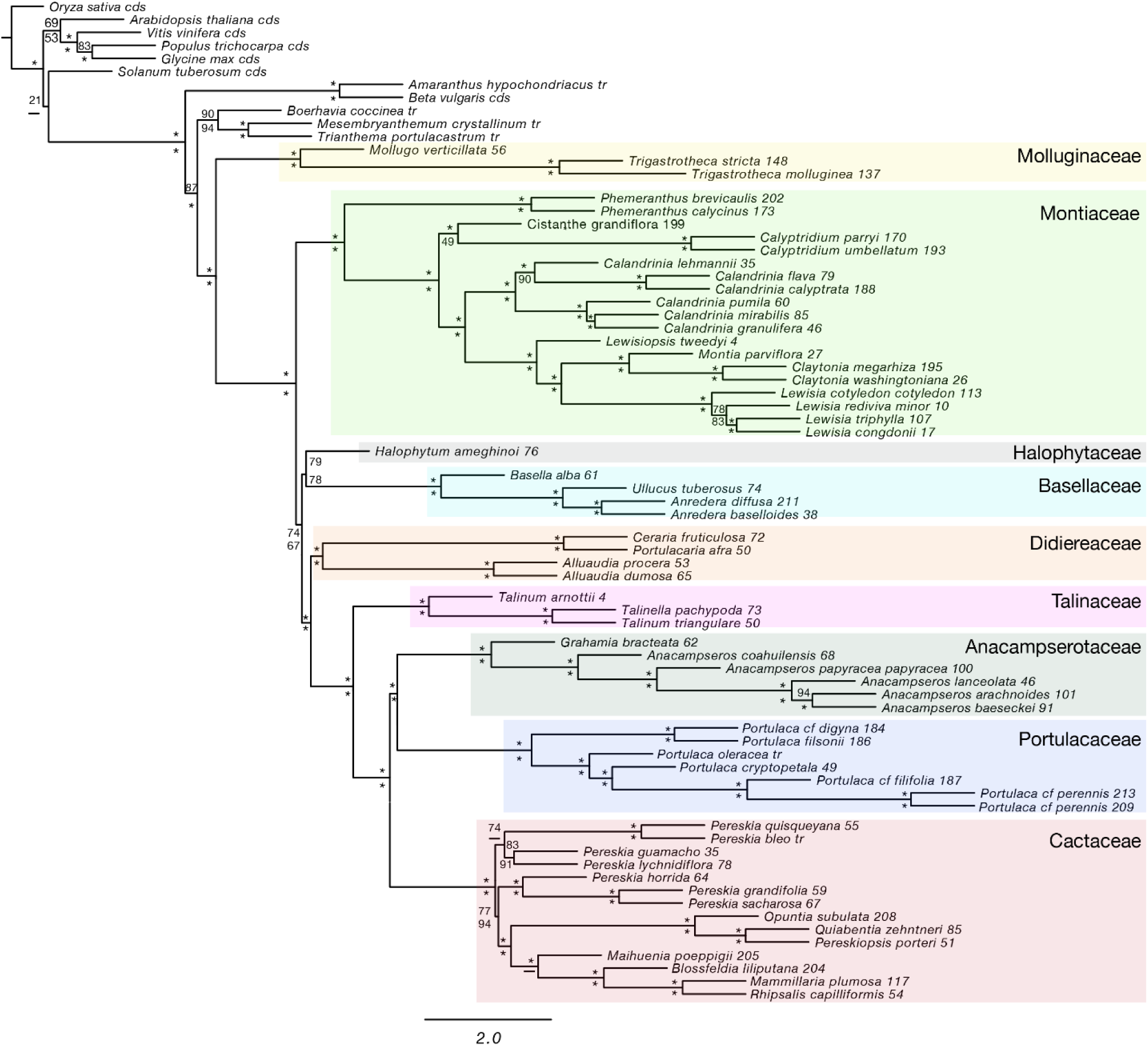
Astral topology from the g5 locus sampling (including only the loci that are present in five or more groups). ASTRAL bootstrap values are above the branches, while RAxML bootstrap values are below the branches. Star indicates greater than 95% bootstrap support.

The major conflict between analyses resided within Cactaceae. While all analyses recover the “core cacti” (sensu Edwards et al. 2005) as monophyletic, four of the five concatenated analyses show *Maihuenia* to be sister to Opuntioideae + Cactoideae with high support, while all ASTRAL analyses and the i57 concatenated analysis recover Opuntioideae as sister to *Maihuenia* plus Cactoideae. The relationships within *Pereskia* are quite variable and are generally poorly supported. Two analyses recover a monophyletic *Pereskia*, two recover *P. lychnidiflora* alone as sister to the core cacti, and the remaining six recover a clade composed of *P. grandifolia, P. sacharosa*, and *P. horrida* as sister to the core cacti, a relationship first proposed by Edwards et al. (2005; the “caulocacti”).

Even though the various species tree analyses demonstrated congruence in the resolution of major relationships, Bayesian Concordance Analysis highlighted significant underlying genome-wide conflict among loci. First, the primary concordance tree from the portullugo-wide analysis revealed similar topologies as the ASTRAL and concatenation analyses, but with low to medium genome-wide concordance factors (CF; from 0.63 for ACP to 0.28 for Portulacineae except Montiaceae; Table S6), indicating that significant portions of the genome support relationships that deviate from the dominant signal (Fig. 4). For instance, in the ACPT analysis, the sister group of Cactaceae as Portulacaceae + Anacampserotaceae was supported by half of our sampled loci (mean CF 0.52), while the other half supported either Anacampserotaceae (mean CF 0.25) or Portulacaceae (mean CF 0.23) alone as sister to Cactaceae (Fig. 4). In the Portulacineae-wide analysis, the ASTRAL and concatenation-inferred position of Halophytaceae as sister to Basellaceae received a mean CF of 0.35, somewhat higher than an alternative placement as sister to Montiaceae (mean CF 0.19; Fig. S4).

**Figure 4.**
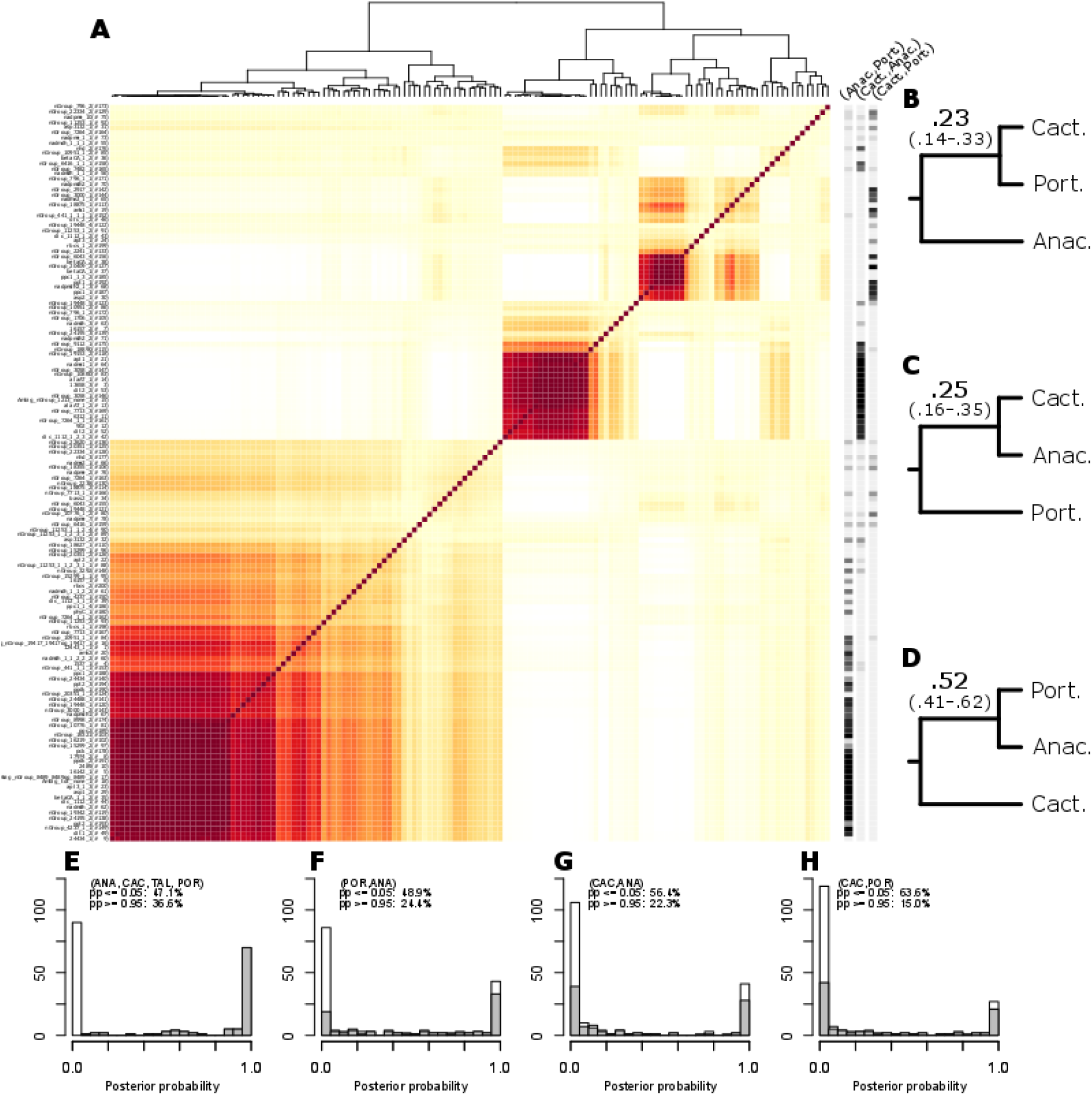
Genomic conflict for phylogenetic relationships between Cactaceae and its putative sister groups, Portulacaceae and Anacampserotaceae. (A) Heatmap of the “calculate-pairs” analysis in BUCKy (i.e., based on a posterior distribution of trees randomly pruned to one exemplar for each of Cactaceae, Portulacaceae, Anacampserotaceae, and Talinaceae), indicating the posterior probability (pp) that a pair of loci support the same topology (red: pp=1; white: pp=0). Thus, each row and column represents a locus; locus names and numbers are given to the left, posterior probability from the MrBayes analysis of individual loci (i.e., based on unpruned trees) for three alternative sister group relations are given to the right (light grey: pp=0; black: pp=1), and a dendrogram based on Euclidean distance between pp values is drawn above. (B-D) Topologies for the three putative sister relations, with genome-wide concordance factor (bold) and 95% credibility interval (in brackets) indicated. (E-H) Histograms of posterior probability from MrBayes analyses of individual loci for four putative clades (E: Anacampserotaceae + Cactaceae + Portulacaceae + Talinaceae; F: Portulacaceae + Anacampserotaceae; G: Cactaceae + Anacampserotaceae; H: Cactaceae + Portulacaceae). Contribution to total frequency by loci in which Anacampserotaceae + Cactaceae + Portulacaceae + Talinaceae is not supported (pp=0) is drawn in white, contribution by other loci is drawn in gray.[Note that strongly supported conflict across genes, rather than lack of information in individual genes, is indicated through presence of multiple major red blocks in panel A, considerable concordance factors for conflicting resolutions in panels B-D; and bimodal distributions in histograms of panels E-H.]

### Molecular Evolution of PEP Carboxylase

One of the major *ppc* paralogs, *ppc-1E1*, has undergone multiple rounds of duplication in ancestral Portulacineae, with sequences clustering into five main paralogs, denoted *ppc1E1a–e* (following Christin et al. 2014; Fig. 5). In addition, *ppc-1E1a* underwent a further duplication (*ppc-1E1a’*) in ancestral *Portulaca*. Members of Didiereaceae and *Lewisia* appear to possess additional copies of *ppc-1E1* distinct from the *ppc-1E1a–e* duplications, though their placement is poorly supported. Non-Portulacineae Caryophyllales possess a single copy of *ppc-1E1*.

**Figure 5.**
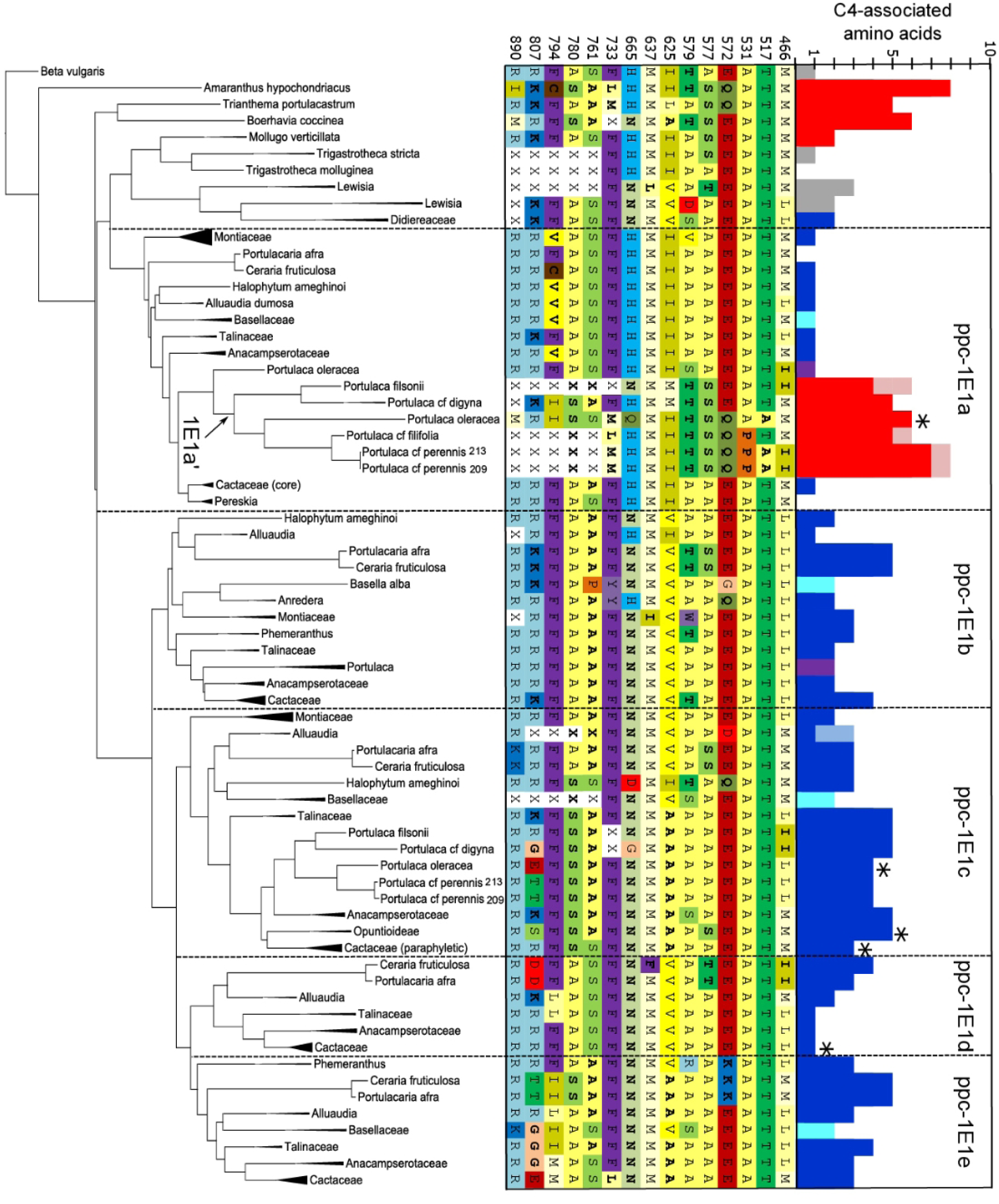
Molecular evolution of *ppc-1E1* in portullugo. Phylogenetic tree of *ppc-1E1* obtained via RAxML. Where possible, named lineages are compressed. Amino acids residues for 14 positions shown to be under positive selection in C_4_ grasses, as well as position 890, which is associated with malate sensitivity, are shown (numbering corresponds to *Zea mays* CAA33317). For compressed lineages, the most frequent amino acid is shown. Amino acids are color-coded based on chemical properties. For specific residues that could not be identified due to missing data or ambiguity, amino acids with highest marginal probabilities are shown, and corresponding color codes are partially transparent (*rerootingMethod* function in phytools, Revell 2012). Amino acids specifically associated with C_4_ in grasses are in boldface, and the number of boldface amino acids for a lineage are indicated with red, blue, gray, and purple horizontal bars. Red bars indicate a C_4_ lineage, blue bars indicate a lineage with CAM activity, light blue bars indicate suspected CAM activity, purple indicates both C_4_ and CAM, and gray indicates a C_3_ lineage. For *Portulaca*, a C_4_ and CAM lineage, *ppc-1E1a’* bars are coded red because of the known association with C_4_ photosynthesis, and *ppc-1E1c* bars are coded blue because of the documented association with CAM activity. Asterisks indicate lineages with drought-induced night time up-regulation of transcript copy number, suggesting relevance to CAM activity.

We inferred multiple C_4_-associated amino acid substitutions in *ppc-1E1*, both inside and outside of Portulacineae. In particular, *ppc-1E1a’* within the *Portulaca* lineage, which has been previously shown to be associated with C_4_ activity (Christin et al. 2014), contains an elevated number of C_4_-associated amino acids relative to *ppc-1E1a* across Portulacineae. We also looked for C_4_-specific AA substitutions in CAM species, with the hypothesis that there may be convergence in coding sequences between these two syndromes due to their shared function of PEP Carboxylase. We discovered a number of C_4_-adaptive osubstitutions in *ppc-1E1b–e* in different CAM lineages, with most lineages showing the greatest accumulation (five) in *ppc-1E1c*. However, some species in particular (*Ceraria* + *Portulacaria*) show a very broad distribution of putative C_4_-like substitutions across *ppc-1E1b–e*. The most ubiquitous and consistent C_4_-adaptive AA in other plant groups, Ser780, has appeared only in *ppc-1E1c* and *ppc-1E1e* (and the C_4_ ppc-1E1a’). Sequences of *ppc-1E2* and *ppc2* were also examined, and both paralogs exhibit very low rates of evolution in general, and a low number of C_4_-associated amino acid substitutions (0.59 and 0.29 C_4_-associated substitutions per site per unit branch length, respectively) relative to *ppc-1E1a–e* (1.44, 1.12, 1.47, 2.37, and 2.45, respectively).

## Discussion

### Targeted Gene Enrichment with Multi-Gene Families

Hybrid bait enrichment is becoming increasingly common in phylogenetics, with researchers developing specialized probe sets designed specifically for unique lineages, much in the way we have done with portullugo (e.g., Lemmon et al. 2012; Nicholls et al. 2015; Heyduk et al. 2016). The critical difference is that most studies focus on capturing ‘single-copy’ genes in plant genomes (De Smet et al. 2013; Chamala et al. 2015), which facilitate contig assembly and homology assignment. Despite the obvious practical advantages of single-copy loci for phylogenetic inference, many questions require sequencing other parts of the genome. By broadening sampling to include multi-copy gene families, targeted enrichment provides an effective means to collect data for phylogenetic reconstruction while simultaneously accumulating comparative datasets on particular gene families of interest. In this study, we attempted to target a wide array of genes in our probe design, and included gene families of major photosynthesis proteins that are relevant to our broader research program.

While all phylogenomic datasets undoubtedly contain some errors in orthology assignment, overall, we feel confident that we are accurately sorting paralogs into their correct ortholog groups, resulting in accurate phylogenetic inference. First, our phylogenetic findings are congruent with those recovered in phylogenetic studies that used a non-controversial, sanger-sequencing-based approach (e.g., the ACPT clade, Portulacineae ex Montiaceae; Nyffeler & Eggli 2010). Secondly, for classic recalcitrant nodes, our BUCKy analyses reveal significant underlying genomic conflict, despite an emerging phylogenetic resolution (discussed below). Thus, rather than conflicting previous studies, our results are largely consistent with them, and provide greater insight into the real genomic conflict underlying historically recalcitrant nodes.

Overall, we have identified two key challenges to working with multi-gene families in hybrid enrichment approaches. First, if baits are designed only from exons (e.g., RNA-seq data, which are the most likely genomic resource for most groups), we can only sequence across short introns. Thus, long introns in the genomic data prevent the entire gene from assembling into one contig, which increases the chance of assembling chimeric sequences derived from multiple paralogs in later steps. Second, the sampling density of the gene family ‘backbone tree’ used to assign orthology has an enormous effect on the ability to accurately classify contigs. Both of these limitations were mostly (though not entirely) overcome by our iteration of the short-read classification step, as confidently placed contigs were maintained in the backbone tree for further rounds of homology assignment. We admit that our approach is largely one of “brute force” at this point, with massive iteration and refinement of key steps. However, as researchers continue to sequence additional taxa with their designed baits, the backbone trees in this step will become more densely sampled, and confidence in contig classification should increase.

### Strong Consensus for Major Relationships within the Portullugo, and the Sister Lineage of the Cacti

The primary goal of our targeted enrichment study was phylogenetic inference, and our analyses provide robust support for most major relationships within the portullugo. In nearly all cases, concatenation and coalescent-based inference methods are congruent and show similar levels of support. Although previous analyses presented conflicting support for the branch uniting Portulacineae with Molluginaceae (Arakaki et al. 2011; Soltis et al. 2011; Yang et al. 2015; Brockington et al. 2015; Thulin et al. 2016), our analyses across all datasets confidently support this node, though our sampling outside of the portullugo is sparse and not designed to directly address this question. Montiaceae consistently appears as sister to the remaining Portulacineae, though with lower support in both ASTRAL and concatenated analyses than we would have predicted. Long recognized clades, like ACPT (Anacampserotaceae, Cactaceae, Talinaceae, Portulacaceae) and ACP (ACPT without Talinaceae) remain strongly supported. More importantly, relationships among other difficult taxa are beginning to crystallize. Our analyses confirm the monophyly of the Didiereaceae *s.l.* (Bruyans et al, 2014) and its placement as sister to the ACPT clade. In addition, Halophytaceae, a monotypic subshrub endemic to the arid interior of Argentina, is now placed with moderate support as sister to Basellaceae in all of our analyses, which is a new finding.

One of the more complex phylogenetic problems in the Portulacineae has been identifying the sister lineage of Cactaceae. The cacti are among the most spectacular desert plant radiations, with ~ 1500 species of mostly stem succulents that diversified recently, during the late Miocene-Pliocene time period (Arakaki et al. 2011). They are closely related to *Portulaca*, a globally widespread, herbaceous and succulent C_4_ lineage, and the Anacampserotaceae, another unusual succulent lineage with most species endemic to South Africa. The relationship among these three clades has remained uncertain, despite numerous phylogenetic studies aimed at resolving it (Hershkovitz and Zimmer 2000; Applequist and Wallace 2001; Nyffeler and Eggli 2010; Ocampo and Columbus 2010; Ogburn and Edwards 2015). We present strong support for *Portulaca* + Anacampserotaceae together as the sister lineage of the cacti, in both concatenation (100% BS) and coalescent (98%) analyses.

In spite of this congruence, our BUCKy analyses revealed strong and significant discord among loci for these relationships, with roughly half of our sampled genome (mean CF 0.52) supporting ((A,P),C) and roughly 25% supporting either (A(P,C)) or (P(A,C)) (Fig. 4; abbreviating lineages by their first letter). It is important to note that this discord among individual gene trees is not derived from poorly supported topologies of individual loci; on the contrary, posterior probabilities for the alternative topologies are routinely very high, mostly with 100% support (Fig 4, panels E–H). Due to the congruence and overall strong support for ((A,P)C) by multiple inference methods and alternative matrices, we tentatively accept this topology and present it as the best working hypothesis for ACP relationship. Nevertheless, we find the amount and strength of conflicting signal throughout the genome quite remarkable. The reconstruction of a single, bifurcating species tree has generally been seen as the ultimate goal of phylogenetics and lack of resolution is typically regarded as a problem that will be solved with the addition of more or better data. However, in a growing number of cases, additional data have only shown the problem to be more complicated, and strong conflict in genome-scale data appears to be the rule, rather than the exception (Scally et al. 2012; Suh et al. 2015; Pease et al. 2016; Brown and Thomson 2016; Shen et al. 2017).

### Genomic Conflict in Deep Time Phylogenetics

Commonly proposed reasons for the existence of recalcitrant nodes in phylogeny reconstruction, beyond lack of phylogenetic information, include homoplasy, incomplete lineage sorting, incorrect homology assignment due to gene duplication and loss, protracted gene flow, and hybridization. Homoplasy was long the preferred explanation for lack of resolution due to conflict (as exemplified by long-branch attraction and the Felsenstein zone), when it was assumed that, in general, gene trees would be congruent with the species tree. Newer data are showing that, while some degree of incomplete lineage sorting (ILS) would always be expected, ILS may be a reasonable explanation for recalcitrant nodes in some instances (Oliver 2013; Suh et al. 2015; Hahn and Nakhleh 2015). Strongly supported incongruence of our various gene trees could be evidence for widespread incomplete lineage sorting at several nodes in our phylogeny, including the split between Anacampserotaceae, Cactaceae, and Portulacaceae, and the relationship of the various species of *Pereskia* to the remainder of the Cactaceae.

The adoption of coalescent theory to resolve ancient nodes has been quickly accepted (e.g., Edwards et al. 2007; Mirarab et al. 2014), though not without some skepticism (Gatesy and Springer 2013; Springer and Gatesy 2014; Gatesy and Springer 2014). Clearly, there is obvious value in evaluating gene trees independently of one another, as they may represent distinct evolutionary histories. Concatenation of very large matrices has also been shown to cause inflated support values, masking significant phylogenetic conflict in the underlying data (Salichos and Rokas 2013). Under the coalescent, the expected degree of deviation of gene trees from the species tree depends on effective ancestral population sizes and generation times (Degnan & Rosenberg 2009). In a series of simulations, Oliver (2013) provided some estimate of ancestral population sizes and generation times needed for the signal of ILS to be recovered in practise; while not impossible, our intuition is that these conditions are not often met in plants, at least in our study system.

We cannot help but consider the diffuse and significant numbers of inferred gene duplications in our dataset (Fig. S3), including many potential losses of paralogs in certain groups. It is true that inference of both paralog presence *and* absence is compromised in any genome sub-sampling approach (hybrid enrichment, RNA-seq, etc.), because the absence of a particular paralog could simply be because the paralog was not captured in the sub-sampling or, in the case of transcriptomes, expressed in the collected tissue. Nevertheless, the ubiquitous and phylogenetically dispersed pattern of our inferred duplications across the portullugo (Fig. S3) implies that, regardless of where precisely these duplications are located, isolated duplications are common along the vast majority of reconstructed branches, and not confined to occasional WGD events. Considering estimated genome-wide rates of gene duplication and loss in other groups (Lynch and Conery 2000; Liu et al. 2014), we wonder if the ILS signal in some of these deep-time phylogenetic studies may be better considered as the “incomplete sorting” of paralogs due to differential paralog fixation following gene duplication and subsequent speciation, rather than a persistent signal of incomplete sorting of alleles alone. In datasets like ours, which span deep nodes and typically include no measure of intraspecific sequence variation, we find it difficult to distinguish between these two scenarios when accounting for gene tree-species tree incongruence.

### Molecular Evolution of PEP Carboxylase

A secondary goal of our study was to design an enrichment scheme that would allow us to simultaneously build a large database of genes relevant to the evolution of C_4_ and CAM photosynthesis. Our previous work on PEPC evolution in this lineage identified five Portulacineae-specific gene duplications within *ppc-1E1*, the major *ppc* paralog that is most often recruited into C_4_ function across eudicots (These duplications all appeared to take place after the separation of the Molluginaceae; Christin et al. 2014; 2015). We also previously identified specific amino acid substitutions in C_4_ and CAM *ppc* loci consistent with changes seen in C_4_ origins in grasses, suggesting that there may be shared adaptive AA residues associated with both C_4_ and CAM function, likely due to the enzyme’s similar function in both syndromes (Christin et al. 2014; 2015). Our small analysis presented here (Fig. 5) is preliminary, and only meant to illustrate the feasibility of performing large-scale comparative molecular evolution studies with bait sequence data by focusing on an already well known gene family as a proof of concept.

Our expanded baits sampling and analysis is consistent with our previous findings. We confirmed the additional duplication of *ppc-1E1a* within the *Portulaca* lineage (*ppc-1E1a’*) that was associated with the evolution of C_4_ photosynthesis in this group, and the use of this specific paralog in C_4_ function has already been documented (Christin et al. 2014). Multiple residues of *ppc-1E1a*’ overlap with amino acids associated with C_4_ photosynthesis in grasses, whereas *ppc-1E1a* possesses 0–1 of the C_4_-associated AA residues in all taxa examined. Strikingly, *ppc-1E1a*’ sequences also exhibit substantial variation within the major clades of *Portulaca*, suggesting that differing C_4_ origins in *Portulaca* were associated with the fixation of distinct C_4_-adaptive AA residues within *ppc-1E1a’* (Christin et al. 2014).

Less is known about the relationship between CAM function and the molecular evolution of PEPC-coding genes. We have discovered the Ser780 residue, which is ubiquitous in C_4_ PEPC in multiple CAM species (Christin et al. 2014); however, in orchids, CAM-expressed PEPC does not seem to require Ser780 (Silvera et al. 2014). Furthermore, expression studies have found nighttime up-regulation of primarily *ppc-1E1c* (with Ser780; Christin et al. 2014; Brilhaus et al. 2015) and in one case each *ppc-1E1a* (Brilhaus et al. 2015), *ppc-1E1d* (Christin et al. 2014), and *ppc-1E1e* (Brilhaus et al. 2015), all without Ser780, in multiple Portulacineae engaged in a CAM cycle. In this first broader look at amino acid substitutions across the entire *ppc* gene family, we can observe a few patterns. First, many CAM species appear to have accumulated multiple AA residues that have been identified as important to C_4_ function, suggesting that there may be a shared selection pressure for both syndromes at the molecular sequence level. Second, only two of the five *ppc1* copies (with the exception of the *Portulaca*-specific *1E1a*’) in Portulacineae have acquired a Ser780: *ppc-1E1c* and *ppc-1E1e*. In general, *ppc-1E1c* and *ppc-1E1e* are the paralogs that have acquired the most C_4_-adaptive AA residues.

A peculiar case is presented by *Ceraria fruticulosa* and *Portulacaria afra* in the Didiereaceae (the sister group to the ACPT clade). These species demonstrate a relatively high number of C_4_-associated amino acids in *ppc-1E1b, ppc-1E1c, ppc-1E1d*, and *ppc-1E1e*; furthermore, the specific residues that overlap with C_4_-associated amino acids largely differ across paralogs, and the only copy in these species with a Ser780 is *ppc-1E1e*. Considering that a *ppc* copy with a Ser780 has, to our knowledge, never been found in non-C_4_ or non-CAM plants, we predict that *ppc-1E1e* in these taxa is primarily used for CAM function. However, the three additional paralogs that also exhibit putatively adaptive AA residues may also contribute to CAM function. This type of scenario has never been demonstrated for C_4_ photosynthesis, though we have previously documented significant upregulation of both *ppc-1E1c* and -*1E1d* in *Nopalea* (CAM, Cactaceae) at night (Christin et al. 2015) and Brilhaus et al. (2015) documented significant upregulation of *ppc-1E1c, -1E1e*, and likely -*1E1a* (*Talinum triangulare*, facultative CAM, Talinaceae).

In light of the broad distribution of putative adaptive residues across the multiple copies of *ppc-1E1* in the Portulacineae, it seems that this lineage and gene family might be an especially powerful system for examining the dynamics of gene duplication and subsequent sub-functionalization (e.g. Ohno 1970). Perhaps in some Portulacineae lineages, functional specialization of particular *ppc-1E1* paralogs took a considerable amount of time post-duplication, with many of them co-contributing to CAM function for millions of years, while accumulating adaptive AA changes independently. Transcriptome profiling of a broader array of Portulacineae could provide critical insight here; for instance, all members of the ACPT clade so far investigated show strong upregulation of *ppc-1E1c* during CAM, potentially because of the presence of the Ser780 residue. Perhaps this mutation occurred earlier in the ACPT clade than it did in the *Ceraria/Portulacaria* clade, which facilitated a more rapid functional specialization of *ppc-1E1c* to CAM function. If the Ser780 mutation is of large effect, then the timing of its appearance may have significant consequences for subsequent specialization of duplicated genes.

In conclusion, we show that it is possible to use targeted sequence capture to sequence gene families across a broad taxonomic range of plants. Phylogenetic studies need not be confined to single copy genes that may be of limited interest outside of their phylogenetic utility; rather, our sampling can be expanded to include large, multi-gene families. Not only does this allow the use of a greater proportion of the genome in targeted sequence capture studies, it also enables exhaustive sampling and analysis of any gene with relevance to a very broad range of evolutionary questions. This creates exciting opportunities for phylogenetic biology in general, opening the potential for systematics-centered research to fully grow into integrative and comprehensive analyses of whole-organism evolution.

## Acknowledgement

The authors would like to thank P.-A. Christin, M. Howison, M. Moeglein, C. Munro, and F. Zapata for helpful discussion; E. Johnson for the figures; B. Dewenter, J.A.M. Holtum, E. van Jaarsveld, F. Obbens, D. Tribble, and R. de Vos for field assistance; B. Dewenter and C. Schorl for lab assistance; the 1KP project for Molluginaceae transcriptome sequences; CapeNature (0028-AAA008-00140), SANParks, and the Government of South Australia (E26345-1) for permission to collect. L.P.H. was supported in part by NSF IGERT grant DGE-0966060.

## Funding

This work was supported by the National Science Foundation (DEB-1252901 to E.J.E.).

## Supplementary Material

Tree files, concatenated alignments, and separate alignments for each locus are available from the Dryad Digital Repository: http://dx.doi.org/10.5061/dryad.[NNNN] (supplementary tables)

## References

Altschul S. F., Gish W., Miller W., Myers E. W., Lipman D. J. 1990. Basic local alignment search tool. J. Mol. Biol. 215:403–410.

Álvarez I., Wendel J.F. 2003. Ribosomal ITS sequences and plant phylogenetic inference. Molecular Phylogenetics and Evolution. 29:417–434.

Ané C., Larget B., Baum D.A., Smith S.D., Rokas A. 2006. Bayesian Estimation of Concordance among Gene Trees. Mol. Biol. Evol. 24:412–426.

Applequist W.L., Wallace R.S. 2001. Phylogeny of the portulacaceous cohort based on ndhF sequence data. Syst. Bot. 206: 406–419.

Arakaki M., Christin P.A., Nyffeler R., Lendel A., Eggli U., Ogburn R.M., Spriggs E., Moore M.J., Edwards E.J. 2011. Contemporaneous and recent radiations of the world’s major succulent plant lineages. Proc. Natl. Acad. Sci. 108:8379–8384.

Baum D.A. 2007. Concordance Trees, Concordance Factors, and the Exploration of Reticulate Genealogy. Taxon. 56:417–426.

Berger S. A., Krompass D., Stamatakis A. 2011. Performance, accuracy, and web server for evolutionary placement of short sequence reads under maximum likelihood. Syst. Biol. 60:291–302.

Besnard G., Muasya A.M., Russier F., Roalson E.H., Salamin N., Christin P.-A. 2009. Phylogenomics of C4 photosynthesis in sedges (Cyperaceae): multiple appearances and genetic convergence. Mol. Biol. Evol. 26:1909–1919.

Brilhaus D., Bräutigam A., Mettler-Altmann T., Winter K., Weber A.P.M. 2015. Reversible Burst of Transcriptional Changes during Induction of Crassulacean Acid Metabolism in *Talinum triangulare*. Plant Physiology. 170:102–122.

Brockington S.F., Yang Y., Gandia Herrero F., Covshoff S., Hibberd J.M., Sage R.F., Wong G.K.S., Moore M.J., Smith S.A. 2015. Lineage-specific gene radiations underlie the evolution of novel betalain pigmentation in Caryophyllales. New Phytol. 207:1170–1180.

Brown J.M., Thomson R.C. 2016. Bayes factors unmask highly variable information content, bias, and extreme influence in phylogenomic analyses. Syst. Biol.:101–14.

Bruyns P.V., Oliveira-Neto M., Melo-de-Pinna G.F., Klak C. 2014. Phylogenetic relationships in the Didiereaceae with special reference to subfamily Portulacarioideae. Taxon. 63:1053–1064.

Chamala S., García N., Godden G.T., Krishnakumar V., Jordon-Thaden I.E., De Smet R., Barbazuk W.B., Soltis D.E., Soltis P.S. 2015. MarkerMiner 1.0: A new application for phylogenetic marker development using angiosperm transcriptomes. Appl. Plant Sci. 3: 1400115.

Chen K., Durand D., Farach-Colton M. 2000. NOTUNG: a program for dating gene duplications and optimizing gene family trees. J. Comp. Biol. 7:429–447.

Christin P.-A., Salamin N., Savolainen V., Duvall M.R., Besnard G. 2007. C4 Photosynthesis Evolved in Grasses via Parallel Adaptive Genetic Changes. Curr. Biol. 17:1241–1247.

Christin P.-A., Arakaki M., Osborne C.P., Bräutigam A., Sage R.F., Hibberd J.M., Kelly S., Covshoff S., Wong G.K.-S., Hancock L., Edwards E.J. 2014. Shared origins of a key enzyme during the evolution of C4 and CAM metabolism. J. Exp. Bot. 65:3609–3621.

Christin P.-A., Arakaki M., Osborne C.P., Edwards E.J. 2015. Genetic enablers underlying the clustered evolutionary origins of C4 photosynthesis in angiosperms. Mol. Biol. Evol. 32:846–858.

Crawford D. J. 2010. Progenitor-derivative species pairs and plant speciation. Taxon 59:1413–1423.

De Bodt S., Maere S., Van de Peer Y. 2005. Genome duplication and origin of Angiosperms. Trends Ecol. Evol. 20:591–597.

De Smet R., Adams K.L., Vandepoele K., Van Montagu M.C.E., Maere S., Van de Peer Y. 2013. Convergent gene loss following gene and genome duplications creates single-copy families in flowering plants. Proc. Natl. Acad. Sci. USA. 110:2898–2903.

de Sousa F., Bertrand Y.J.K., Nylinder S., Oxelman B., Eriksson J.S., Pfeil B.E. 2014. Phylogenetic Properties of 50 Nuclear Loci in Medicago (Leguminosae) Generated Using Multiplexed Sequence Capture and Next-Generation Sequencing. PLoS ONE. 9:e109704–10.

Degnan J.H., Rosenberg, N.A., 2009. Gene tree discordance, phylogenetic inference and the multispecies coalescent. Trends in Ecol. Evol. 24:332–340.

Dohm J. C., Minoche A. E., Holtgräwe D., Capella-Gutiérrez S., Zakrzewski F., Tafer H., Rupp O., Sörensen T. R., Stracke R., Reinhardt R., Goesmann A., Kraft T., Schulz B., Stadler P. F., Schmidt T., Gabaldón T., Lehrach H., Weisshaar B., Himmelbauer H. 2014. The genome of the recently domesticated crop plant sugar beet (*Beta vulgaris*). Nature. 505:546–549.

Dunn C.W., Hejnol A., Matus D.Q., Pang K., Browne W.E., Smith S.A., Seaver E., Rouse G.W., Obst M., Edgecombe G.D., Sørensen M.V., Haddock S.H.D., Schmidt-Rhaesa A., Okusu A., Kristensen R.M., Wheeler W.C., Martindale M.Q., Giribet G. 2008. Broad phylogenomic sampling improves resolution of the animal tree of life. Nature. 452:745–749.

Edgar R. C. 2004. MUSCLE: multiple sequence alignment with high accuracy and high throughput. Nucl. Aci. Res. 32:1792–1797.

Edwards E.J., Nyffeler R., Donoghue M.J. 2005. Basal cactus phylogeny: Implications of Pereskia (Cactaceae) paraphyly for the transition to the cactus life form. Am. J. Bot. 92:1177–1188.

Edwards E.J., Ogburn R.M. 2012. Angiosperm Responses to a Low-CO2 World: CAM and C4 Photosynthesis as Parallel Evolutionary Trajectories. Int. J. Plant Sci. 173:724–733.

Edwards S.V., Liu L., Pearl D.K. 2007. High-resolution species trees without concatenation. Proc. Natl. Acad. Sci. USA. 104:5936–5941.

Faircloth B.C., McCormack J.E., Crawford N.G., Harvey M.G., Brumfield R.T., Glenn T.C. 2012. Ultraconserved elements anchor thousands of genetic markers spanning multiple evolutionary timescales. Syst. Biol. 61:717–726.

Freeling M. 2009. Bias in plant gene content following different sorts of duplication: tandem, whole-genome, segmental, or by transposition. Annu. Rev. Plant Biol. 60:433–453.

Gatesy J., Springer M.S. 2013. Concatenation versus coalescence versus “concatalescence.” Proc. Natl. Acad. Sci. USA. 110:1179–1179.

Gatesy J., Springer M.S. 2014. Phylogenetic analysis at deep timescales: Unreliable gene trees, bypassed hidden support, and the coalescence/concatalescence conundrum. Mol. Biol. Evol. 80:231–266.

Hahn M.W., Nakhleh L. 2015. Irrational exuberance for resolved species trees. Evolution. 70:7–17.

Hershkovitz M.A., Zimmer E.A. 2000. Ribosomal DNA evidence and disjunctions of western American Portulacaceae. Mol. Phyl. Evol. 15:419–439.

Heyduk K., McKain M.R., Lalani F., Leebens-Mack J. 2016. Evolution of a CAM anatomy predates the origins of Crassulacean acid metabolism in the Agavoideae (Asparagaceae). Mol. Phyl. Evol. 105:102–113.

Hudspeth, R. L., & Grula, J. W. 1989. Structure and expression of the maize gene encoding the phosphoenolpyruvate carboxylase isozyme involved in C4 photosynthesis. Plant Mol. Biol. 12:579–589.

Jiao Y., Wickett N.J., Ayyampalayam S., Chanderbali A.S., Landherr L., Ralph P.E., Tomsho L.P., Hu Y., Liang H., Soltis P.S., Soltis D.E., Clifton S.W., Schlarbaum S.E., Schuster S.C., Ma H., Leebens-Mack J., dePamphilis C.W. 2011. Ancestral polyploidy in seed plants and angiosperms. Nature. 473:97–100.

Katoh K., Standley D. M. 2013. MAFFT multiple sequence alignment software version 7: improvements in performance and usability. Mol. Biol. Evol. 30:772–780.

Kersey P.J., Allen J.E., Armean I., Boddu S., Bolt B.J., Carvalho-Silva D., Christensen M., Davis P., Falin L.J., Grabmueller C., Humphrey J., Kerhornou A., Khobova J., Aranganathan N.K., Langridge N., Lowy E., McDowall M.D., Maheswari U., Nuhn M., Ong C.K., Overduin B., Paulini M., Pedro H., Perry E., Spudich G., Tapanari E., Walts B., Williams G., Tello Ruiz M., Stein J., Wei S., Ware D., Bolser D.M., Howe K.L., Kulesha E., Lawson D., Maslen G., Staines D.M. 2016. Ensembl Genomes 2016: more genomes, more complexity. Nucleic Acids Res. 44:D574–80.

Kircher M., Sawyer S., Meyer, M. 2012. Double indexing overcomes inaccuracies in multiplex sequencing on the Illumina platform. Nucl. Aci. Res. 40:e3.

Larget B.R., Kotha S.K., Dewey C.N., Ane C. 2010. BUCKy: Gene tree/species tree reconciliation with Bayesian concordance analysis. Bioinformatics. 26:2910–2911.

Leaché A.D., Rannala B. 2011. The accuracy of species tree estimation under simulation: a comparison of methods. Syst. Biol. 60:126–137.

Lemmon A.R., Emme S.A., Lemmon E.M. 2012. Anchored hybrid enrichment for massively high-throughput phylogenomics. Syst. Biol. 61:727–744.

Li C., Hofreiter M., Straube N., Corrigan S., Naylor G.J.P. 2013. Capturing protein-coding genes across highly divergent species. BioTechniques 54:321–326.

Liu S., Liu Y., Yang X., Tong C., Edwards D., Parkin I.A.P., Zhao M., Ma J., Yu J., Huang S., Wang X., Wang J., Lu K., Fang Z., Bancroft I., Yang T.-J., Hu Q., Wang X., Yue Z., Li H., Yang L., Wu J., Zhou Q., Wang W., King G.J., Pires J.C., Lu C., Wu Z., Sampath P., Wang Z., Guo H., Pan S., Yang L., Min J., Zhang D., Jin D., Li W., Belcram H., Tu J., Guan M., Qi C., Du D., Li J., Jiang L., Batley J., Sharpe A.G., Park B.-S., Ruperao P., Cheng F., Waminal N.E., Huang Y., Dong C., Wang L., Li J., Hu Z., Zhuang M., Huang Y., Huang J., Shi J., Mei D., Liu J., Lee T.-H., Wang J., Jin H., Li Z., Li X., Zhang J., Xiao L., Zhou Y., Liu Z., Liu X., Qin R., Tang X., Liu W., Wang Y., Zhang Y., Lee J., Kim H.H., Denoeud F., Xu X., Liang X., Hua W., Wang X., Wang J., Chalhoub B., Paterson A.H. 1AD. The Brassica oleracea genome reveals the asymmetrical evolution of polyploid genomes. Nat Commun. 5:1–11.

Lynch M., Conery J.S. 2000. The evolutionary fate and consequences of duplicate genes. Science. 290:1151–1155.

Mandel J.R., Dikow R.B., Funk V.A. 2015. Using phylogenomics to resolve mega-families: An example from Compositae. J. Syst. Evol. 53:391–402.

Martin A.P., Burg T.M. 2002. Perils of Paralogy: Using HSP70 Genes for Inferring Organismal Phylogenies. Syst. Biol. 51:570–587.

Matasci N., Hung L.-H., Yan Z., Carpenter E.J., Wickett N.J., Mirarab S., Nguyen N., Warnow T., Ayyampalayam S., Barker M., Burleigh J.G., Gitzendanner M.A., Wafula E., Der J.P., dePamphilis C.W., Roure B., Philippe H., Ruhfel B.R., Miles N.W., Graham S.W., Mathews S., Surek B., Melkonian M., Soltis D.E., Soltis P.S., Rothfels C., Pokorny L., Shaw J.A., DeGironimo L., Stevenson D.W., Villarreal J.C., Chen T., Kutchan T.M., Rolf M., Baucom R.S., Deyholos M.K., Samudrala R., Tian Z., Wu X., Sun X., Zhang Y., Wang J., Leebens-Mack J., Wong G.K.-S. 2014. Data access for the 1,000 Plants (1KP) project. GigaSci. 3:17–10.

McCormack J.E., Hird S.M., Zellmer A.J., Carstens B.C., Brumfield R.T. 2013. Applications of next-generation sequencing to phylogeography and phylogenetics. Molecular Phylogenetics and Evolution. 66:526–538.

Mirarab S., Reaz R., Bayzid M.S., Zimmermann T., Swenson M.S., Warnow T. 2014. ASTRAL: genome-scale coalescent-based species tree estimation. Bioinformatics. 30:i541–8.

Nicholls J.A., Pennington R.T., Koenen E.J.M., Hughes C.E., Hearn J., Bunnefeld L., Dexter K.G., Stone G.N., Kidner C.A. 2015. Using targeted enrichment of nuclear genes to increase phylogenetic resolution in the neotropical rain forest genus Inga (Leguminosae: Mimosoideae). Front. Plant Sci. 6:1382–1420.

Novikova P. Y., Hohmann N., Nizhynska V., Tsuchimatsu T., Ali J., Muir G., Guggisberg A., Paape T., Schmid K., Fedorenko O. M., Holm S., Säll T., Schlötterer C., Marhold K., Widmer A., Sese J., Shimizu K. K., Weigel D., Krämer U., Koch M. A., Nordborg M. 2016. Sequencing of the genus *Arabidopsis* identifies a complex history of nonbifurcating speciation and abundant trans-specific polymorphism. Nature Genetics 48:1077–1082.

Nyffeler R., Eggli U. 2010. Disintegrating Portulacaceae: A new familial classification of the suborder Portulacineae (Caryophyllales) based on molecular and morphological data. Taxon. 59:227–240.

Ocampo G., Columbus J.T. 2010. Molecular phylogenetics of suborder Cactineae (Caryophyllales), including insights into photosynthetic diversification and historical biogeography. Am. J. Bot. 97:1827–1847.

Ogburn M.R., Edwards E.J. 2015. Life history lability underlies rapid climate niche evolution in the angiosperm clade Montiaceae. Mol. Phyl. Evol. 92:181–192.

Ohno S. 1970. Evolution by gene duplication. London: George Alien & Unwin Ltd. Berlin, Heidelberg and New York: Springer-Verlag

Oliver J.C. 2013. Microevolutionary processes generate phylogenomic discordance at ancient divergences. Evolution. 67:1823–1830.

Paulus J.K., Schlieper D., Groth G. 2013. Greater efficiency of photosynthetic carbon fixation due to single amino-acid substitution. Nat. Commun. 4:1518.

Pease J.B., Haak D.C., Hahn M.W., Moyle L.C. 2016. Phylogenomics Reveals Three Sources of Adaptive Variation during a Rapid Radiation. PLoS Biol. 14:e1002379–24.

R Core Team. 2016. R: A language and environment for statistical computing. R Foundation for Statistical Computing, Vienna, Austria. https://www.R-project.org/.

Renny-Byfield S., Wendel J.F. 2014. Doubling down on genomes: Polyploidy and crop plants. Am. J. Bot. 101:1711–1725.

Revell L.J. 2012. phytools: an R package for phylogenetic comparative biology (and other things). Methods Ecol. Evol. 3:217–223.

Ronquist F., Teslenko M., van der Mark P., Ayres D.L., Darling A., Höhna S., Larget B., Liu L., Suchard M.A., Huelsenbeck J.P. 2012. MrBayes 3.2: efficient Bayesian phylogenetic inference and model choice across a large model space. Syst. Biol. 61:539–542.

Salichos L., Rokas A. 2013. Inferring ancient divergences requires genes with strong phylogenetic signals. Nature. 497:327–331.

Scally A., Dutheil J.Y., Hillier L.W., Jordan G.E., Goodhead I., Herrero J., Hobolth A., Lappalainen T., Mailund T., Marques-Bonet T., McCarthy S., Montgomery S.H., Schwalie P.C., Tang Y.A., Ward M.C., Xue Y., Yngvadottir B., Alkan C., Andersen L.N., Ayub Q., Ball E.V., Beal K., Bradley B.J., Chen Y., Clee C.M., Fitzgerald S., Graves T.A., Gu Y., Heath P., Heger A., Karakoc E., Kolb-Kokocinski A., Laird G.K., Lunter G., Meader S., Mort M., Mullikin J.C., Munch K., O’Connor T.D., Phillips A.D., Prado-Martinez J., Rogers A.S., Sajjadian S., Schmidt D., Shaw K., Simpson J.T., Stenson P.D., Turner D.J., Vigilant L., Vilella A.J., Whitener W., Zhu B., Cooper D.N., de Jong P., Dermitzakis E.T., Eichler E.E., Flicek P., Goldman N., Mundy N.I., Ning Z., Odom D.T., Ponting C.P., Quail M.A., Ryder O.A., Searle S.M., Warren W.C., Wilson R.K., Schierup M.H., Rogers J., Tyler-Smith C., Durbin R. 2012. Insights into hominid evolution from the gorilla genome sequence. Nature. 483:169–175.

Schmickl R., Liston A., Zeisek V., Oberlander K., Weitemier K., Straub S.C.K., Cronn R.C., Dreyer L.L., Suda J. 2016. Phylogenetic marker development for target enrichment from transcriptome and genome skim data: the pipeline and its application in southern African Oxalis (Oxalidaceae). Mol. Ecol. Resour. 16:1124–1135.

Shen X.-X., Hittinger C.T., Rokas A. Contentious relationships in phylogenomic studies can be driven by a handful of genes. Nature Ecology & Evolution. 1:0126 EP –.

Silvera K., Winter K., Rodriguez B.L., Albion R.L., Cushman J.C. 2014. Multiple isoforms of phosphoenolpyruvate carboxylase in the Orchidaceae (subtribe Oncidiinae): implications for the evolution of crassulacean acid metabolism. J. Exp. Bot. 65:3623–3636.

Soltis P.S., Marchant D.B., Van de Peer Y., Soltis D.E. 2015. Polyploidy and genome evolution in plants. Curr. Opin. Genet. Dev. 35:119–125.

Springer M.S., Gatesy J. 2014. Land plant origins and coalescence confusion. Trends Plant Sci. 19:267–269.

Stamatakis A. 2014. RAxML version 8: a tool for phylogenetic analysis and post-analysis of large phylogenies. Bioinformatics 30:1312–1313.

Stolzer M., Lai H., Xu M., Sathaye D., Vernot B., Durand D. 2012. Inferring duplications, losses, transfers and incomplete lineage sorting with nonbinary species trees. Bioinformatics 28:409–415.

Suh A., Smeds L., Ellegren H. 2015. The Dynamics of Incomplete Lineage Sorting across the Ancient Adaptive Radiation of Neoavian Birds. PLoS Biol. 13:e1002224.

Thulin M., Moore A.J., El-Seedi H., Larsson A., Christin P.-A., Edwards E.J. 2016. Phylogeny and generic delimitation in Molluginaceae, new pigment data in Caryophyllales, and the new family Corbichoniaceae. Taxon. 65:775–793.

Wickett N.J., Mirarab S., Nguyen N., Warnow T., Carpenter E., Matasci N., Ayyampalayam S., Barker M.S., Burleigh J.G., Gitzendanner M.A., Ruhfel B.R., Wafula E., Der J.P., Graham S.W., Mathews S., Melkonian M., Soltis D.E., Soltis P.S., Miles N.W., Rothfels C.J., Pokorny L., Shaw A.J., DeGironimo L., Stevenson D.W., Surek B., Villarreal J.C., Roure B., Philippe H., dePamphilis C.W., Chen T., Deyholos M.K., Baucom R.S., Kutchan T.M., Augustin M.M., Wang J., Zhang Y., Tian Z., Yan Z., Wu X., Sun X., Wong G.K.-S., Leebens-Mack J. 2014. Phylotranscriptomic analysis of the origin and early diversification of land plants. Proc. Natl. Acad. Sci. USA 111:E4859–E4868.

Yang Y., Moore M.J., Brockington S.F., Soltis D.E., Wong G.K.-S., Carpenter E.J., Zhang Y., Chen L., Yan Z., Xie Y., Sage R.F., Covshoff S., Hibberd J.M., Nelson M.N., Smith S.A. 2015. Dissecting molecular evolution in the highly diverse plant clade Caryophyllales using transcriptome sequencing. Mol. Biol. Evol. 32: 2001–2014

